# Risperidone regulates the expression of schizophrenia-related genes in the murine forebrain

**DOI:** 10.1101/2025.08.18.670814

**Authors:** Magdalena Ziemiańska, Mateusz Zięba, Anna Radlicka-Borysewska, Łukasz Szumiec, Sławomir Gołda, Małgorzata Borczyk, Marcin Piechota, Michał Korostyński, Jan Rodriguez Parkitna

## Abstract

Risperidone acts on monoaminergic signaling to alleviate psychosis. At the cellular level, through both direct and indirect effects, the drug induces a specific pattern of gene expression, which differs between the basal ganglia and frontal cortex. These risperidone-regulated changes influence neuronal plasticity and are crucial for both its antipsychotic and extrapyramidal effects. Here, we employed sequencing-based spatial transcriptomics to comprehensively characterize gene expression changes in the male mouse (*Mus musculus* L.) forebrain after an acute dose of risperidone (0.5 mg/kg, i.p.). The transcriptional patterns were structure-specific, and unsupervised clustering of spatial profiles accurately identified cortical divisions, layers, and basal ganglia subregions. Differential gene expression was subsequently analyzed within each anatomically defined cluster using a customized statistical framework. Risperidone significantly altered the levels of 95 transcripts across 12 brain regions. The largest number of changes was observed in ventral brain areas, including the olfactory tubercle (25 differentially regulated transcripts), the diagonal band nucleus (22), the corpus callosum and commissures (13), and the lateral septal nucleus (9). Notably, 21 of the 95 differentially expressed genes were previously associated with schizophrenia, including *Olig2*, *Smpd3*, and *Cacna1i*. Overall, our results indicate that the strongest effects of risperidone are in medial and ventral brain regions rich in oligodendrocytes and glial cells. Furthermore, the enrichment analysis provides robust evidence of a molecular link between the drug’s mechanism of action and genetic factors involved in schizophrenia.

**Highlights:**

- Unsupervised clustering accurately identifies transcripts’ localization in the brain
- Acute risperidone treatment alters spatial transcriptional patterns of 95 genes
- Risperidone-regulated transcripts include 21 genes previously linked to schizophrenia
- Spatial gene expression analysis offers novel insight into the drug action mechanism

## 1. Introduction

Antipsychotic drugs act on monoaminergic signaling to diminish the positive symptoms of schizophrenia (Kane, 1987). While they exert their effects primarily through dopamine D2 receptor blockade, second-generation antipsychotics are also potent antagonists of serotonin 5-HT2A receptors and interact with various monoaminergic and cholinergic targets (Meltzer, 2013; Miyamoto et al., 2005; Seeman, 2002). A high relative ratio of 5-HT2A and D2 receptor antagonism defines atypical antipsychotics, which cause fewer adverse extrapyramidal effects while still effectively reducing psychosis (Meltzer, 2013). It is important to note that the dopaminergic mechanism of action of antipsychotic drugs is a major piece of evidence supporting the dopamine hypothesis of schizophrenia’s etiology (Howes et al., 2015, 2017; Snyder et al., 1974). Nonetheless, despite progress in psychosis pharmacotherapy, the root causes of schizophrenia remain unclear, treatments are limited in addressing negative symptoms, and ongoing research into how antipsychotics work continues to reveal increasing complexity.

A key approach to unraveling these molecular mechanisms is to profile drug effects on gene expression (Harrison and Weinberger, 2005; Nakamura and Takata, 2023; Robertson et al., 1994). Changes in transcription support long-term plasticity, so drug-induced gene expression profiling offers insight into neuronal mechanisms behind both immediate and lasting effects of psychotropic drugs (McClung and Nestler, 2008; Zygmunt et al., 2018b). Antipsychotic treatment has been shown to robustly induce expression of activity-related transcripts, especially immediate early genes (de Bartolomeis et al., 2017; Dragunow et al., 1990; Robertson and Fibiger, 1992). The distribution of drug-induced transcriptional changes varies by region, mainly affecting the nucleus accumbens and frontal cortex, with differences between drugs in regional expression patterns (de Bartolomeis et al., 2017). It has been observed that typical antipsychotics exert stronger effects on activity-regulated genes in the dorsal striatum. In contrast, atypical antipsychotics, which tend to cause fewer extrapyramidal adverse effects, show relatively greater effects in the prefrontal cortex and nucleus accumbens (Deutch and Duman, 1996; Robertson et al., 1994; Wan et al., 1995). At the same time, dopaminergic dysfunction in schizophrenia exhibits a distinct spatial pattern, with hyperactivity in the basal ganglia and hypoactivity in the prefrontal cortex being the prevailing view (Guillin et al., 2007; Tzschentke, 2001). Therefore, the spatial pattern of antipsychotic effects is essential for their effectiveness and may also predict the severity of extrapyramidal side effects.

Early studies on transcriptional responses to psychotropic drugs concentrated on selected immediate-early genes such as Fos. The use of microarrays (Fehér et al., 2005; Ikeda et al., 2010; Korostynski et al., 2013) and later RNA-sequencing technologies (Crespo-Facorro et al., 2015; Zygmunt et al., 2018a) expanded this approach to transcriptome-wide analyses, allowing unbiased detection of drug effects on gene expression. These studies identified specific cellular and molecular pathways involved in pharmacological treatments, such as the neuroplasticity-related MAPK signaling pathway (Korostynski et al., 2013) or calcium signaling (Fehér et al., 2005). Because of technical limitations, these approaches usually focus on individual brain structures or neuronal populations, leaving the overall spatial organization of transcriptional responses across the brain only partly understood. Additionally, the gene expression changes caused by these drugs are often nonspecific: similar transcriptomic responses are observed after treatment with both antipsychotics and psychostimulants (Zygmunt et al., 2018b). Although these changes occur in different cell populations (Bertran-Gonzalez et al., 2008), they become indistinguishable when tissue is homogenized. Furthermore, most psychotropic drugs activate glucocorticoid receptor-dependent transcription in glial cells, probably reflecting adaptations to altered neuronal activity (Zygmunt et al., 2018b). Recently, the first single-cell RNA-seq analysis of gene expression induced by antipsychotics in the striatum identified highly drug-specific gene patterns and revealed transcriptional changes in microglia (Abrantes et al., 2022). Therefore, while single-cell analysis offers unprecedented insight into the molecular effects of antipsychotics, the method remains limited to specific brain areas due to technical constraints.

Here we perform a high-resolution spatial analysis of risperidone-induced gene expression changes in the murine forebrain to assess the effects of an antipsychotic within the brain’s native tissue architecture (Ståhl et al., 2016). We have selected risperidone as an atypical antipsychotic extensively used in the treatment of psychoses, schizophrenia, and bipolar disorder (Chopko and Lindsley, 2018). Our results show that risperidone induces spatially discrete transcriptional changes that affect the expression of genes previously linked to schizophrenia through genome-wide association studies.

## 2. Methods

### 2.1 Animals

Experiments were performed on adult male mice (10 weeks old C57BL/6N mice, Charles River Laboratories, Germany) weighing 20–26 g. Mice were housed 4 per cage at the animal facility of the Maj Institute of Pharmacology of the Polish Academy of Sciences under standard conditions (22 °C, 12/12 h light/dark cycle) with free access to food and water. All animal experiments were planned following the ARRIVE guidelines (Sert et al., 2020), and performed following the European and Polish law (Directive 2010/63/UE, European Convention for the Protection of Vertebrate Animals Used for Experimental and other Scientific Purposes ETS No.123, and Polish Law Dz.U. 2015 poz. 266). All procedures were approved by the II Local Institutional Animal Care and Use Committee in Kraków (permit no. 136/2022).

### 2.2 Drug treatment and tissue preparation

Risperidone was suspended in a 1% v/v Tween 80 solution (Sigma-Aldrich, St. Louis, MO, USA) in saline by sonication. Mice were habituated to intraperitoneal injections with saline for two consecutive days before receiving a single acute injection of risperidone (0.5 mg/kg in a volume of 5 μl/g body weight; Tocris, 2865) or vehicle. Animals were sacrificed 2 h after risperidone treatment, brains were collected, rinsed with cooled saline (4 °C), and coated in OCT compound (Cell Path, UK). Then, they were snap-frozen in Peel-A-Way® embedding molds (Polysciences, USA) by immersion in isopentane chilled with dry ice (-78 °C). Samples were stored at -80 °C in sealed containers for long-term preservation. Before slicing, brains embedded in the OCT compound were equilibrated to the cryostat chamber temperature (-20 °C) for at least 30 minutes. Tissue blocks were trimmed, and 10 μm-thick coronal sections were cut using a CM 3050S cryostat microtome (Leica Microsystems, Germany) at an object temperature of -13 °C and a chamber temperature of -20 °C. For spatial transcriptomics, one coronal section containing the prefrontal cortex and rostral striatum (1.54–1.18 mm from bregma) was collected from each animal. Coronal sections were mounted one per capture area on a pre-cooled (-20 °C) Visium Spatial slide. Sections were fixed in methanol at -20 °C for 30 minutes and stained with hematoxylin and eosin according to the manufacturer’s protocol (Tissue Preparation Guide Rev. D; Methanol Fixation, H&E Staining & Imaging for Visium Spatial Protocol Rev. D, Visium Spatial Protocols). Imaging was performed using a Leica DMi8 microscope with Leica Application Suite X v3.5.2.18963 software. Each section was imaged in bright field using an HC PL FLUOTAR 10x NA: 0.30 (DRY) lens and a DFC 7000 camera with the following settings: color depth 8 bit, resolution 1920×1440 pixels, gain 10, color saturation ∼22, intensity 21, aperture 24, and exposure time 4.5 ms. Images were saved in the .tif format and used to determine the surface area of the array covered by tissue in Loupe Browser v6.0 (10x Genomics). Processed images were then used to generate .json files for further analysis in the spaceranger-MACS3-Nanopore-spaceranger processing pipeline.

### 2.4 Visium library preparation and sequencing

After imaging, brain slices were enzymatically permeabilized for 8 minutes, followed by incubation in reverse transcription master mix at 53 °C for 45 minutes. The resulting cDNA was incubated with 0.08 M KOH and washed with EB buffer. Next, second-strand synthesis (65 °C, 15 minutes), denaturation (0.08 M KOH, 10 min), and cDNA amplification were performed. The total number of cDNA amplification cycles was determined by qPCR using KAPA SYBR FAST qPCR Master Mix (Roche, Switzerland) with 1 μl of cDNA solution. Amplified cDNA was purified using SPRIselect beads (Beckman Coulter, USA). Quantification and quality control were conducted using Bioanalyzer 2100 (Agilent, Palo Alto, CA, USA) with High Sensitivity DNA Kit reagents (Agilent, Palo Alto, CA, USA) to determine total cDNA yield. Spatial gene expression libraries were constructed according to the manufacturer’s guidelines (Visium Spatial Gene Expression Reagent Kits User Guide Rev G and Visium Spatial Gene Expression Library Construction). First, cDNA samples were fragmented with post-fragmentation end-repair and A-tailing (32 °C for 5 min, 65 °C for 30 s). Next, double-sided size selection with SPRIselect was done before the adaptor ligation and post-ligation cleanup. cDNA amplification with a unique index pair was conducted according to the protocol, followed by a second double-sided size selection with SPRIselect beads. The quality of the libraries was assessed using the Bioanalyzer 2100 with High Sensitivity DNA Kit reagents. Samples were stored at -20 °C for long-term preservation. Pooling and sequencing of the libraries were performed externally at CeGaT (Germany) using the NovaSeq 6000 Sequencing System (SP flow cell, Illumina). Paired-end, dual-indexed RNA sequencing was performed using recommended parameters for Visium Spatial Gene Expression - read 1, 28 cycles; i7 index, 10 cycles; i5 index, 10 cycles; and read 2, 50 cycles, 200 million read pairs per sample.

### 2.5 Sequencing data analysis

RNA sequencing yielded, on average, ∼332 million total reads per library. Raw RNAseq data quality was assessed with FastQC (version 0.11.5-cegat). Samples had an average Q30 score of 92.95%. Read alignment was performed using the Space Ranger v1.3.1 (10x Genomics) pipeline, and then a custom approach was developed in our laboratory. First, the reference genome, tissue section micrographs, and FASTQ files were processed using the ‘spaceranger count’ pipeline (10x Genomics) to detect tissue boundaries, identify fiducial frame markers, and perform barcode/UMI counting. Then, reads were aligned to the mm10 mouse reference genome (GRCm38/mm10; 2020-A; June 23, 2020). The resulting BAM read alignment files were merged into a single experiment-wide file using Samtools (https://github.com/samtools/samtools) and subsequently processed with MACS3 (https://github.com/macs3-project/MACS). RNA-seq reads were mapped to reference genome sequences, and coverage at specific sites was converted to “peaks”. Strand orientation was determined using Samtools, based on coverage depth in both strand directions —a larger proportion of reads in one direction qualified the strand as forward (+) or reverse (-).

Next, peak filtering was performed using the following criteria: peak amplitude > 400, read counts per peak > 1200, count-to-amplitude ratio > 1.4, score value > 350, and summit. p. log> 35. The identified peaks were annotated to gene or LTR sequences. This annotation used a one-sided margin, with peaks located less than 30,000 bp from the gene sequence. Peak coordinates were then compared by intersection (± 30,000 bp from the 5’ end of the transcript) with long-read sequencing data from Oxford Nanopore Technologies, obtained in a previous mouse striatum study (Chrószcz et al., 2025). A total of 15,243 peaks were identified, corresponding to 10,493 protein-coding genes (Sequence Read Archive, accession no PRJNA1143882). A new reference transcriptome was created using ‘spaceranger mkref’, based on the processed and annotated peak coordinates (GitHub - ippas/ifpan-janrod-spatial). The resulting reference and FASTA files were processed with ‘spaceranger count’ to generate feature-barcode matrices quantifying transcript abundance at each tissue spot. After aligning to the reference genome, principal component analysis (PCA) was performed, followed by clustering with Seurat v 4.4.0. A shared nearest neighbor (SNN) graph was constructed from the PCA results and used as input for UMAP visualization. Quantile normalization was applied to each cluster, and the average transcript expression was evaluated using the customized bioinformatics workflow. The identification of structure- and tissue-specific markers was validated with marker combinations reported in literature and databases (see Supplementary Table 1). Clusters with fewer than 20 spots were excluded from statistical comparisons. Mann–Whitney U tests were used on the mean expression values within clusters. Differential gene expression was considered significant if the p-value < 0.01, the log₂ ratio ≥ 0.8, and the relative expression in control and/or experimental samples ≥ 0.2. The false discovery rate (FDR) per cluster was estimated using Monte Carlo permutation tests (12 samples, 1000 iterations) while maintaining a fixed clustering resolution of 0.4.

### 2.6 Ontology analyses

Gene set enrichment analysis of 95 risperidone-regulated genes was performed using the Enrichr web tool (Kuleshov et al., 2016). Results from the GeDiPNet 2023 (Kundu et al., 2023) and GWAS Catalog 2023 (Cerezo et al., 2025) libraries were used to identify significantly overrepresented associations with schizophrenia. Functional relationships among the 21 risperidone-regulated genes associated with schizophrenia were examined using the Enrichr-KG tool (Evangelista et al., 2023). Analyses were conducted with four annotation sets: ChEA3 2022 (transcription factor regulation), MGI Mammalian Phenotype Level 4 2021 (phenotypic annotations), GO Biological Processes 2021 (biological functions), and Reactome 2022 (signaling pathways). These associations were used to build a knowledge graph summarizing enriched biological processes, regulatory networks, and phenotypic links.

### 2.7 Data accessibility

RNAseq datasets are available from the Sequence Read Archive under accession number PRJNA1143882. Bright-field H&E tissue images are available upon request. Scripts used for data analysis are available at: https://github.com/ippas/ifpan-janrod-spatial/tree/master.

## 3. Results

### 3.1 Spatial patterns of gene expression in the forebrain

Drug effects on gene expression were assessed in coronal forebrain sections (1.54–1.18 mm from bregma) from 10-week-old male mice that received an acute i.p. injection of 0.5 mg/kg risperidone or saline 2h before brain collection. Sections were processed to construct 12 cDNA libraries, 6 from animals treated with risperidone and 6 from controls. The average cDNA fragment size was 440 bp. Reads were aligned to the GRCm38 reference genome. To analyze the data, we first performed unsupervised clustering of profiles from all individual spots across all sections (19.473 spots) using Seurat v4.4.0. A resolution parameter of 0.4 was selected, resulting in 17 clusters (#1 to #17), the minimum number needed to recapitulate the neuroanatomical structure of the forebrain sections in our dataset. Cluster sizes ranged from 8 to 307 spots across sections. A UMAP projection of the clustering results is shown in Fig. 1A. Clusters representing the nucleus accumbens (#15), striatum (#17), and olfactory tubercle (#11) are close together, consistent with the common presence of medium spiny neurons in these regions. Conversely, the corpus callosum (#6), characterized by dense fiber tracts and a high content of oligodendrocytes and glia, is located near the diagonal band (#12) and lateral septal nucleus (#13), which also contain many fibers and oligodendrocytes. Clusters representing cortical divisions (#1–4, #7–10, #14) are also grouped in the UMAP projection.

**Fig. 1.**
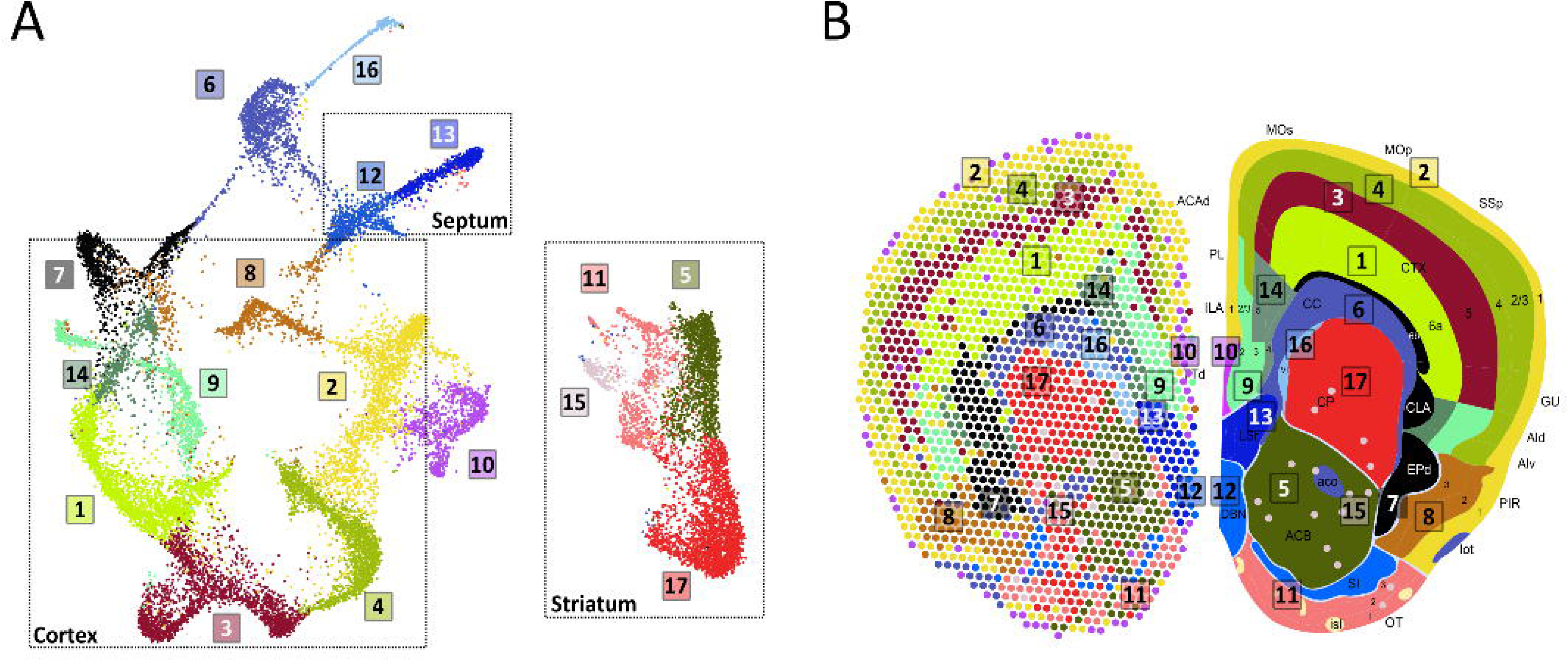
Spatial mapping of structural gene expression in the murine forebrain for region-specific drug effects analysis. A. UMAP projection of unsupervised clustering of gene expression profiles corresponding to all individual spots. Colors represent spots constituting each of the 17 clusters. B. Left: Spatial localization of spots in a representative brain section, with colors depicting cluster identities. Right: Coronal section of the mouse brain adapted from the Allen Reference Atlas – Mouse Brain. 1, 2, 3, 4, 5, 6a, 6b – layers of the cortex, ACAd-anterior cingulate area, dorsal part; ACB – nucleus accumbens; aco – anterior commissure; Ald – agranular insular cortex, dorsal part; Alv - agranular insular cortex, ventral part; CLA - claustrum; CP – caudatoputamen (dorsal striatum); CTX – cerebral cortex, DBN - diagonal band nuclei; EPd – endopiriform nucleus, dorsal part; fa – anterior forceps of the corpus callosum; GU – gustatory cortex; ILA – intralimbic cortex; isl – Islands of Calleja; lot – lateral olfactory tract; LSr – lateral septal nucleus, rostral part; MOp – primary motor cortex; MOs – supplementary motor area; OT – olfactory tubercle; PIR – piriform cortex; PL – prelimbic cortex; SI – substantia innominata; SSp – primary somatosensory cortex; TTd – Taenia tecta, dorsal part; VL – lateral ventricle.

As shown in Fig. 1B, the spatial arrangement of the clusters in a representative section closely matches the anatomical organization of the forebrain. Consequently, clusters are enriched with structure-specific markers described in the literature, supporting their anatomical identity as listed in Supplementary Table 1. Based on their anatomical location and gene expression profiles, clusters #2 and #10 correspond to the brain meninges and cortical layer 1, respectively, characterized by numerous non-neuronal markers (DeSisto et al., 2020; Remsik et al., 2021). Clusters #4, #3, and #1 represent inner cortical layers: cluster #4 aligns with layers 2/3, marked by genes such as *Cux2*, *Kcnip2*, and *Rasgrf2* (Del-Aguila et al., 2019; Maynard et al., 2021); cluster #3 corresponds to layers 4/5, characterized by *Rorb* expression (Del-Aguila et al., 2019); and cluster #1 matches layers 5/6, with markers like *Grik4*, *Sulf2*, and *Rprm* (Del-Aguila et al., 2019; Maynard et al., 2021). Clusters #9 and #14 include the medial prefrontal cortex (prelimbic and infralimbic areas, taenia tecta), insular cortex, and gustatory areas, expressing genes such as *Enc1*, *Fezf2*, and *Mef2c* (Cahoy et al., 2008; Salem et al., 2024; Skene et al., 2018). Cluster #8 likely represents the ventral layers of the piriform cortex, characterized by inhibitory interneuron markers, including *Cck*, *Sst*, and *Ntng1* (based on the interneuron marker list from PanglaoDB (Franzén et al., 2019)). The claustrum, endopiriform nucleus, and cortical layer 6b are grouped within cluster #7, expressing genes such as *Rprm*, *Oprk1*, *Nr4a2*, and *Cbln2* (Del-Aguila et al., 2019). Cluster #6 comprises the corpus callosum, anterior commissure, and lateral olfactory tract, identified by oligodendrocyte markers like *Mog*, *Mbp*, *Opalin*, and *Mobp* (Horiuchi et al., 2017; Skene et al., 2018). Adjacent to the corpus callosum, cluster #16 corresponds to the subventricular zone, indicated by neuronal progenitor markers such as *Igfbp5* and *Sox2* (Lachmann et al., 2018). Additionally, the analysis distinguished between the dorsal (cluster #17) and ventral parts of the striatum (clusters #5 and #11). Cluster #17 matches the caudatoputamen, expressing markers like *Ido1*, *Drd2*, and *Kcnip2* (Skene et al., 2018). Cluster #5 corresponds to the nucleus accumbens, expressing medium spiny neuron markers including *Drd1*, *Drd2*, *Bcl11b*, and *Adora2a*; cluster #15 likely represents dispersed striatal cells expressing *Kcnk2* and *Bcl11* (Puighermanal et al., 2020). Cluster #11 is identified as the olfactory tubercle, based on the expression of genes such as *Otof1*, *Pcp4*, and *Ppp1r2* (list compiled by (Diamant et al., 2025) from the Allen Atlas, (Lein et al., 2007)). Medial to the striatum, cluster #12 corresponds to the diagonal band nucleus, marked by *Lhx6*, *Rab3b*, *Elfn1*, and *Gad2* (Diamant et al., 2025), while the adjacent cluster #13 represents the lateral septal nucleus, enriched in *Trpc4*, *Ecel1*, *Gfra1*, and *Cxcl14* (Diamant et al., 2025).

### 3.2 Risperidone-induced differential gene expression

Differential gene expression caused by acute risperidone treatment (0.5 mg/kg) was assessed within each cluster. Transcript abundance was normalized using quantile normalization across brain sections, and average expression levels were calculated for all spatial spots assigned to each cluster per section. Clusters with fewer than 20 spots were excluded from further analysis. Differential expression was evaluated using the Mann–Whitney U test, with significance defined as a log₂(fold change) of at least ±0.8, a p-value < 0.01, and a minimum mean transcript abundance of 0.2 in at least one group. A total of 95 genes met these criteria, with 76 upregulated and 19 downregulated following risperidone treatment. Four genes — *Ddit4*, *Elk1*, *Sgk1*, and *1110059E24Rik* (C9orf85) — showed regulation in multiple regions, while the remaining 91 genes exhibited significant abundance changes in a single structure. Significant gene expression differences were observed in 12 of 17 clusters (Fig. 2). False discovery rates were calculated separately for each cluster to account for multiple testing and are listed next to the respective clusters in Fig. 2. Most gene expression changes were seen in subcortical areas (Fig. 3). The greatest number of changes was found in the olfactory tubercle (#11) with 25 transcripts regulated by risperidone, including activity-responsive genes such as *Homer1*, *Egr3*, and *Elk1*. Next was the diagonal band nucleus (#12) with 22 risperidone-regulated genes, including *Sorl1* and *Rab11a*. There were 9 differentially expressed transcripts in the adjacent lateral septal nucleus (#13), including *Smpd3* and *Fah2*. A total of 13 genes were upregulated in the brain commissures (#6), including *Ncam1* and *Kcna1*. Several oligodendrocyte markers, such as *Sgk1*, *Olig2*, and *Lpar1*, were also elevated.

**Fig. 2.**
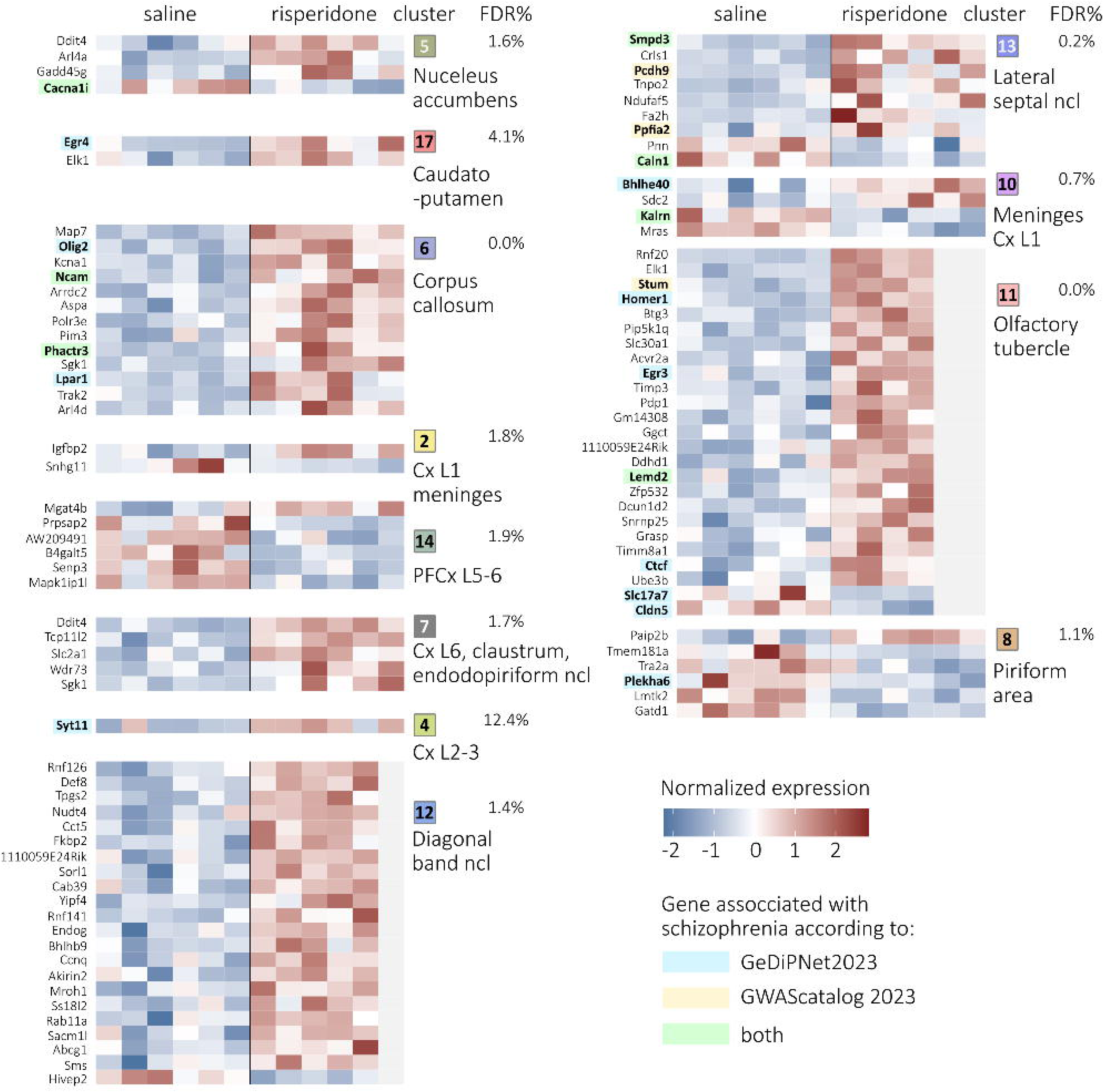
Risperidone-induced gene expression alterations in the murine forebrain. RNA-seq results with spatial localization (left) are presented as heatmaps displaying 95 transcripts with transcriptome-wide significance (FDR permutation-based per cluster). The heatmaps illustrate transcript abundance 2 hours after risperidone injection, with color intensity proportional to the standardized value (z-score between −2 and 2) as indicated on the adjacent color scale. The complete list of 95 drug-affected transcripts is shown on the left. Schizophrenia-associated genes are highlighted in blue - 18 genes enriched for the schizophrenia term in the GeDiPNet 2023 database (p = 1.71 × 10⁻⁴, q-value), yellow - 10 genes identified in human GWAS studies from the GWAS Catalog 2023 (p = 0.02, q = 0.26) and green - the 7 overlapping genes associated with schizophrenia in both GeDiPNet 2023 and GWAS Catalog 2023.

**Fig. 3.**
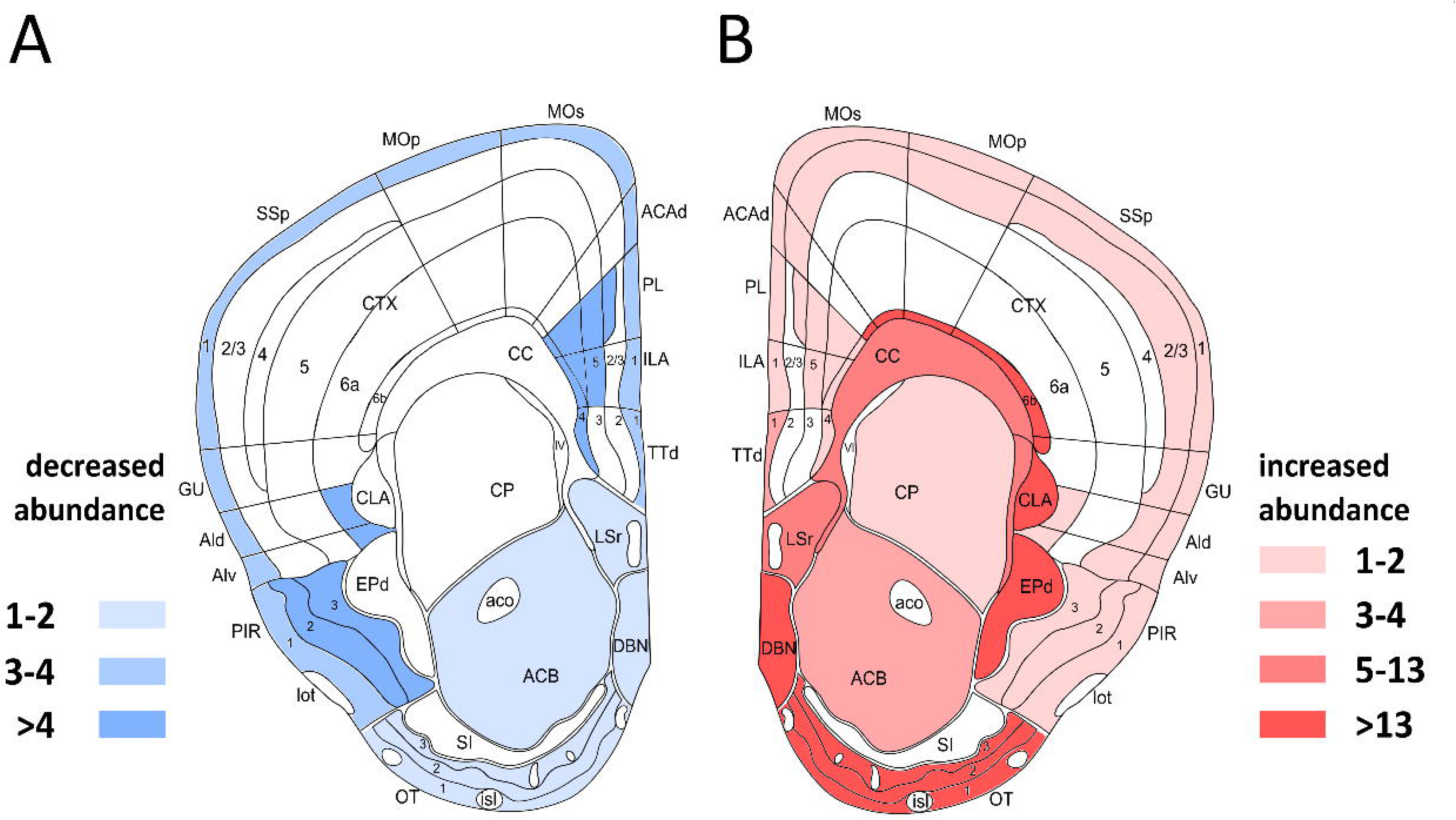
Regional distribution of 95 differentially expressed genes mapped onto a coronal section of the mouse brain, adapted from the Allen Reference Atlas – Mouse Brain. A. The number of downregulated transcripts per brain region. B. The number of upregulated transcripts per brain region. Color intensity reflects the number of transcripts with altered abundance within each cluster.

Taken together, the largest number of expression changes occurred in glia-enriched regions — including the commissures (#6), diagonal band (#12), lateral septal nucleus (#13), and olfactory tubercle (#11)— with most genes showing increased transcript levels (Fig. 3). Conversely, cortical regions such as dorsal cortical layers 5 and 6 (#1), the piriform cortex (#8), and top cortical layers and meninges (#2, #10) primarily showed downregulated gene expression following risperidone exposure. Interestingly, the dorsal and ventral striatum exhibited relatively few significant changes, despite having the highest levels of dopamine receptor D2 expression, which is the main pharmacological target of risperidone. In the dorsal striatum (#17), only two activity-dependent transcription factors, *Egr4* and *Elk1*, were significantly induced, while in the nucleus accumbens (#5), changes in the expression of *Ddit4*, *Arl4a*, *Gadd45g*, and *Cacna1i* were noted.

### 3.3 Enrichment analyses of risperidone-regulated genes

We have performed enrichment analysis on the 95 risperidone-regulated genes using Enrichr and annotations from the Gene Disease Pathway Networks database (“GeDiPNet 2023”, (Kundu et al., 2023)) that integrates data from all major clinical databases (including DisGeNET, ClinGen, ClinVar, HPO, OrphaNet, and PsyGeNET) and the NHGRI-EBI GWAS Catalog, the most comprehensive database of gene-trait associations compiled from genome-wide association (GWAS) studies (“GWAS Catalog 2023”, (Cerezo et al., 2025)). The most significant annotation overrepresentation in the GeDiPNet 2023 database was observed for the association with ‘schizophrenia’, with 18 genes among 95 risperidone-regulated transcripts (Fisher’s exact test, p = 1.71 × 10⁻⁴, q = 0.0568). These included *Egr3*, *Homer1*, *Cldn5*, *Lemd2*, *Slc17a7*, and *Ctcf* in the olfactory tubercle (#11); *Egr4* in the caudatoputamen (#17); *Phactr3*, *Olig2*, *Ncam1*, and *Lpar1* in the corpus callosum (#6); *Plekha6* in the piriform cortex (#8); *Bhlhe40* and *Kalrn* in the meninges/layer 1 of the cortex (#2); *Smpd3* and *Caln1* in the lateral septal nucleus (#13); *Cacna1i* in the nucleus accumbens (#5); and *Syt11* in cortical layers 2 and 3 (#4). Moreover, 10 of the 95 risperidone-regulated genes were annotated as schizophrenia-associated in the Enrichr GWAS Catalog 2023 library — trait “schizophrenia” (MONDO_0005090) (Fisher’s exact test, p = 0.0198, q = 0.264). These genes included *Smpd3, Ppfia2, Caln1 and Pcdh9* (lateral septal nucleus, #13) *Cacna1i,* (nucleus accumbens, #5)*, Phactr3, Ncam1,* (corpus callosum, #6)*, Lemd2, Stum* (olfactory tubercle, #11), and *Kalrn* (meninges and top cortical layers, #10). In total, 21 genes were associated with schizophrenia in the GeDiPNet 2023 database or the GWAS Catalog 2023, with 7 genes listed by both sources. Brain region-specific changes in the expression of risperidone-regulated genes enriched for schizophrenia in both the GeDiPNet 2023 database and the GWAS Catalog 2023 are presented in Fig. 4 (increases in abundance) and Fig. 5 (decreases in abundance). *Phactr3* (Ikeda et al., 2010; Li et al., 2017) and *Ncam1* (Dang et al., 2025; Trubetskoy et al., 2022) were upregulated in the areas corresponding to the commissures (#6, Fig.4), *Smpd3* (Trubetskoy et al., 2022) in the lateral septal nucleus (#13), and *Lemd2* (Trubetskoy et al., 2022) in the olfactory tubercle (#11). In turn, *Cacna1i* (Lam et al., 2019; Li et al., 2017) had reduced expression in the nucleus accumbens (#5, Fig. 5), *Caln1* (Lam et al., 2019; Li et al., 2017) was decreased in the lateral septal nucleus (#13), and *Kalrn* (Ikeda et al., 2010; Li et al., 2017) in the top cortical layers and/or the meninges (#2). Additionally, risperidone-regulated genes reported as associated with schizophrenia in previous human studies, including *Ppfia2* (Levinson et al., 2012), *Pcdh9*, *Stum* (Trubetskoy et al., 2022), and *Cldn5* (Lindsay, 2001) are shown in Supplementary Figure 1.

**Fig. 4.**
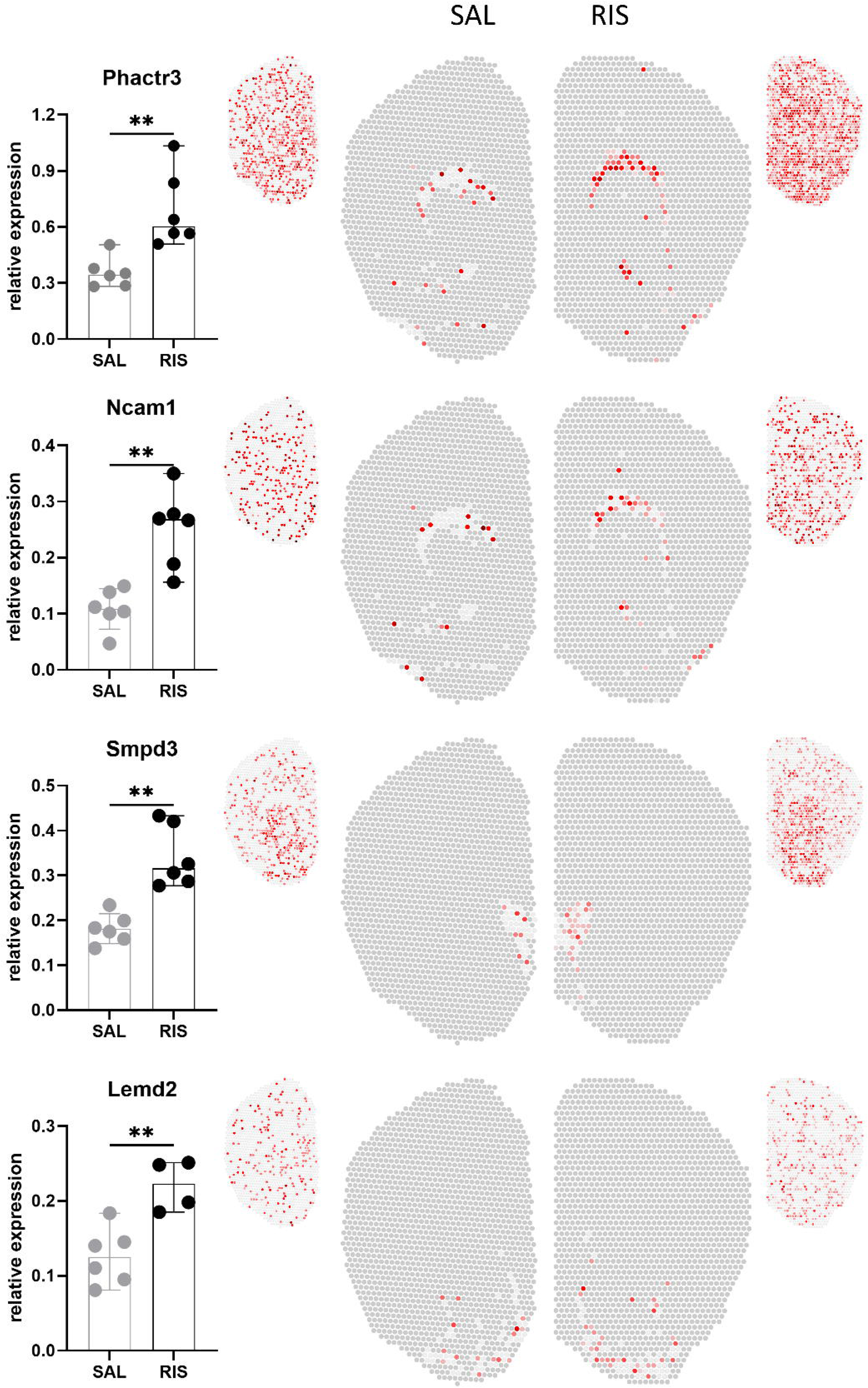
Spatial gene expression profiles of transcripts with increased abundance following acute risperidone treatment. The bar plots (left) and spatial gene expression profiles (right) illustrate the relative expression levels of genes upregulated by risperidone and associated with schizophrenia in both GeDiPNet 2023 and GWAS Catalog 2023. Bar plots show medians with 95% confidence intervals, based on Mann– Whitney U test results (**p1<10.01). Full expression profiles are displayed alongside the main images. Statistically significant changes were observed only in specific clusters, as outlined in the main panels.

**Fig. 5.**
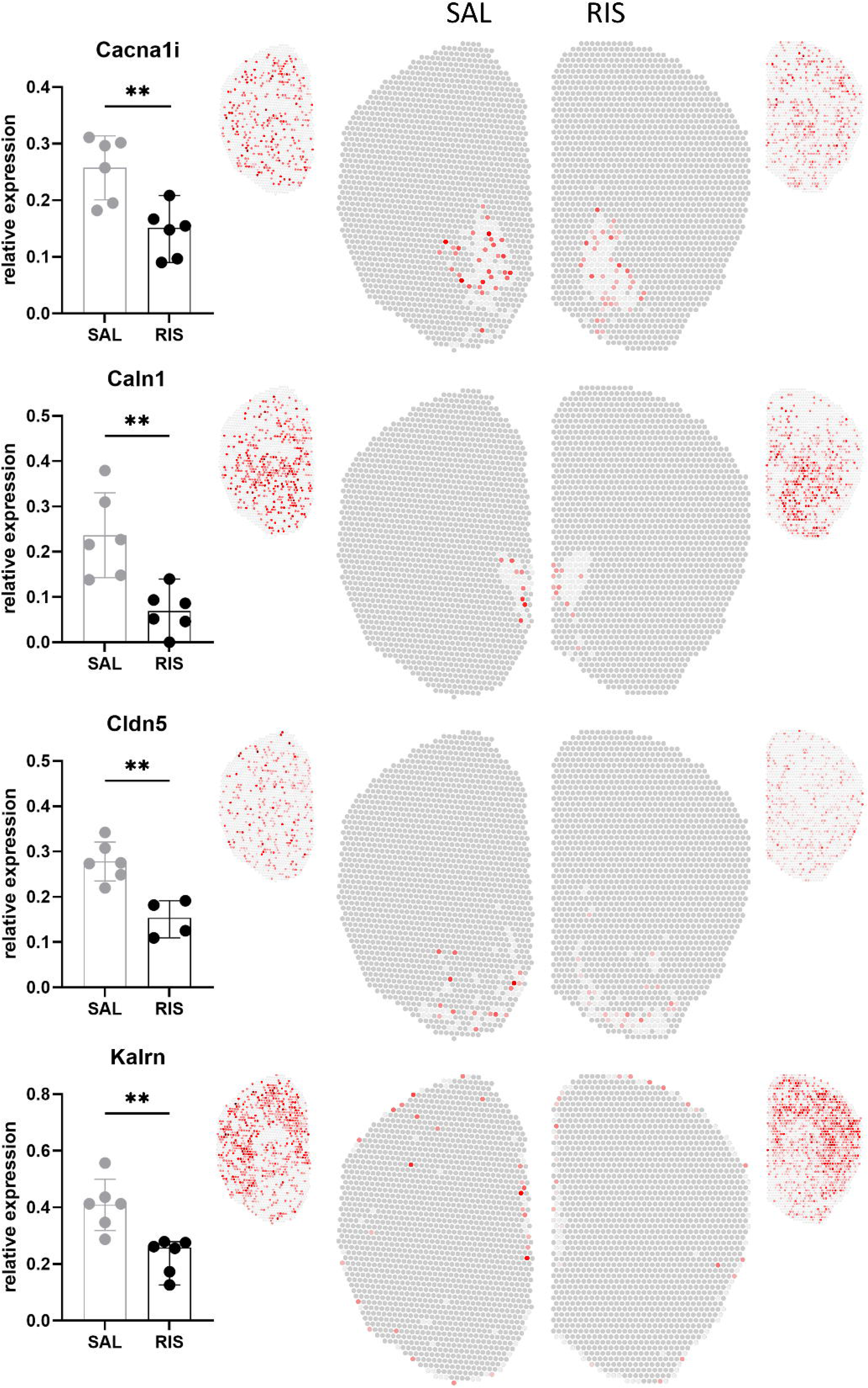
Spatial gene expression profiles of transcripts with decreased abundance following acute risperidone treatment. The bar plots (left) and spatial gene expression profiles (right) illustrate the relative expression levels of genes downregulated by risperidone and associated with schizophrenia in both GeDiPNet 2023 and GWAS Catalog 2023. Bar plots show medians with 95% confidence intervals, based on Mann–Whitney U test results (p1<10.01). Full expression profiles are displayed alongside the main images. Statistically significant changes were observed only in specific clusters, as outlined in the main panels.

**Fig. 6.**
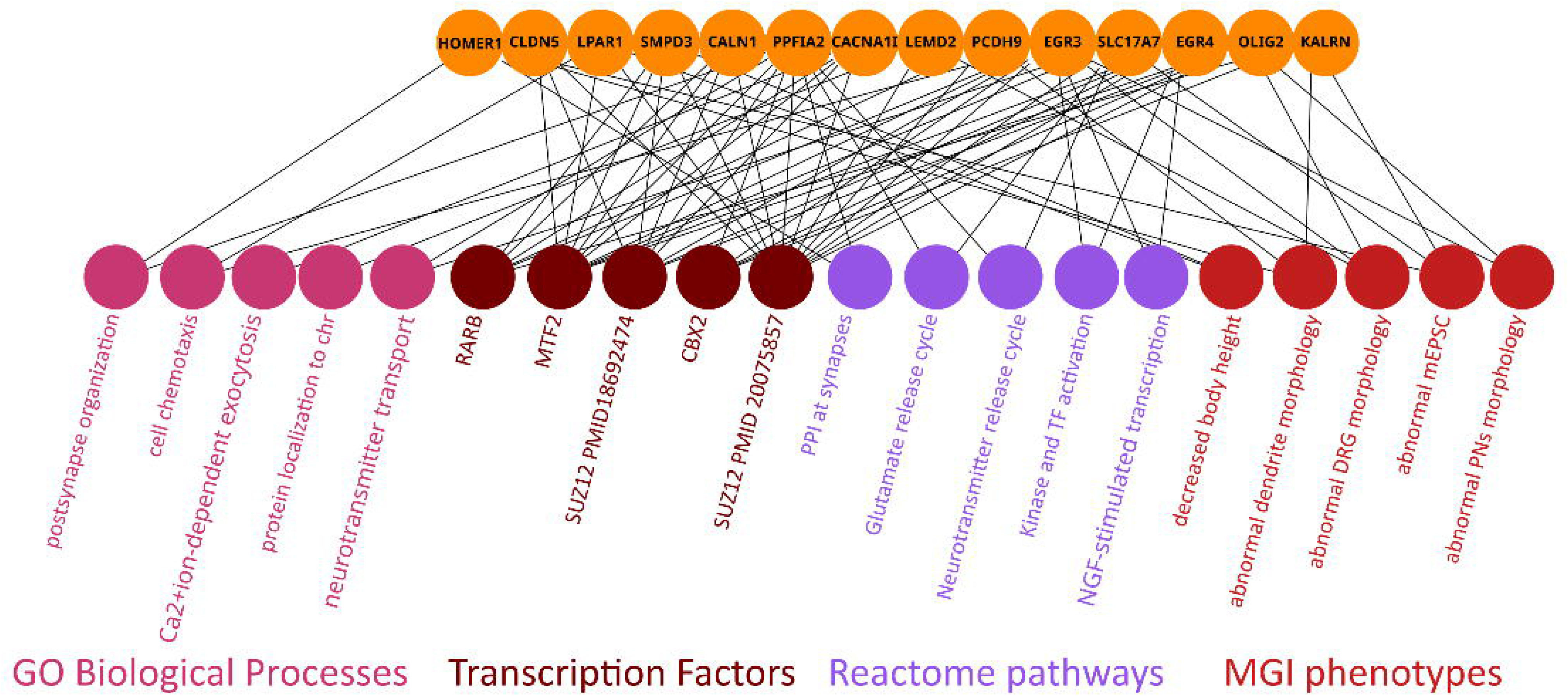
Overview of functional connections among risperidone-regulated genes linked to schizophrenia. The knowledge graph displays associations with enriched categories: transcription factor regulatory elements from ChEA3 2022 (brown), morphological and functional phenotypes from MGI Mammalian Phenotype Level 4 2021 (dark red), biological processes from GO Biological Processes 2021 (pink), and cellular pathways from Reactome 2022 (purple). The graph was prepared based on the results obtained by using the Enrichr-KG tool. The complete list of terms is enclosed in Supplementary Table 3.

Lastly, we reanalyzed ontology enrichment in the subset of 21 genes associated with schizophrenia. Analysis of transcription factor binding sites in gene promoter regions (based on chromatin immunoprecipitation data) showed an overrepresentation of Polycomb group factors, with SUZ12 sites detected in 12 promoters (PMID: 20075857, p = 1.24 × 10⁻⁷, q = 7.04 × 10⁻⁵; PMID: 18692474, p = 1.11 × 10⁻⁵, q = 2.18 × 10⁻³). MTF2 was found in 10 promoters (p = 1.15 × 10⁻⁵, q = 2.18 × 10⁻³), and CBX2 in 5 promoters (p = 2.86 × 10⁻⁵, q = 4.06 × 10⁻³). We also performed analyses for enrichment of ontology terms using GO Biological Processes 2021, Reactome 2022, and MGI Mammalian Phenotype Level 4 2021. Results showed enrichment in Gene Ontology Biological Processes, including, for example, *Ppfia2* and *Slc17a7* in neurotransmitter transport (GO:0006836, p = 2.64 × 10⁻³, q = 9.44 × 10⁻²), Cacna1i and Syt11 in the regulation of calcium ion-dependent exocytosis (GO:0017158, p = 4.794 × 10⁻⁴, q = 4.541 × 10⁻²), and *Homer1* and *Ppfia2* in the regulation of postsynapse organization (GO:0099175, p = 3.904 × 10⁻⁴, q = 4.541 × 10⁻²). Reactome pathway analysis further linked *Ppfia2* and *Slc17a7* to the Glutamate Neurotransmitter Release Cycle (R-HSA-210500, p = 2.62 × 10⁻⁴, q = 2.36 × 10⁻²) and, among others, *Ppfia2* and *Homer1* to Protein–Protein Interactions at Synapses (R-HSA-6794362, p = 3.81 × 10⁻³, q = 6.85 × 10⁻²). Finally, MGI Mammalian Phenotype set analysis identified, for example, *Homer1*, *Slc17a7*, and *Kalrn* as linked to abnormal miniature excitatory postsynaptic currents (MP:0004753, p = 1.34 × 10⁻⁴, q = 9.91 × 10⁻³), and *Homer1*, *Pcdh9*, and *Kalrn* with abnormal dendrite morphology (MP:0008143, p = 5.68 × 10⁻⁵, q = 5.60 × 10⁻³). These findings suggest that schizophrenia-associated genes are not only regulated by common transcription factors, mainly Polycomb group proteins, but also converge functionally in pathways important for synaptic transmission and neuronal structure.

## 4. Discussion

Two key insights emerge from the spatial analysis of drug-regulated spatial gene expression patterns. First, transcriptional responses to risperidone are specific to brain regions, with distinct sets of genes regulated in adjacent structures. Second, there is a notable overrepresentation of genes related to schizophrenia risk among the differentially regulated transcripts, suggesting that antipsychotic treatment influences molecular pathways involved in the disease’s pathophysiology.

We observed strong risperidone-induced changes in transcript levels in subcortical regions of the forebrain, which largely align with prior findings on gene expression changes following antipsychotic treatment in animal models (Chen and Chen, 2005; de Bartolomeis et al., 2017; Fehér et al., 2005; Korostynski et al., 2013; Zygmunt et al., 2018) and post-mortem samples from schizophrenia patients (Merikangas et al., 2022). Several genes that are differentially expressed, including *Bhlhe40*, *Egr3*, *Egr4*, *Elk1*, *Sgk1*, *Ddit4*, and *Homer1a*, are activity-dependent transcripts previously reported to be acutely regulated by antipsychotics in the striatum, cortex, and corpus callosum (Korostynski et al., 2013; Zygmunt et al., 2018). Most of these changes occurred in the septum—areas rich in white matter—as well as the olfactory tubercle and piriform cortex, rather than the nucleus accumbens or prefrontal cortex. Moreover, some well-studied activity-regulated transcripts in the context of antipsychotics, such as *Fos*, *Fosb*, or *Egr1*, are absent from our results. The overall difference in spatial patterns likely stems from the historical focus on preselected brain regions directly involved in dopaminergic pathology of schizophrenia and from findings based on in situ hybridization experiments assessing *Fos* expression. Notably, early studies reported antipsychotic-induced gene expression changes in the olfactory tubercle, septum, and ventral cortical regions (Cools et al., 1995; Robertson and Fibiger, 1992). However, these findings were not followed up with extensive research. The pattern of risperidone-induced gene expression overlaps only slightly with the distribution of dopamine D2 and serotonin 5-HT2A receptors. In the forebrain, D2 receptors are mainly found in a subset of medium spiny neurons within the striatum, nucleus accumbens, and olfactory tubercle, as well as in the upper layers of the cortex (Cieslak et al., 2024; Gangarossa et al., 2013; Meador-Woodruff et al., 1989). Conversely, 5-HT2A receptors are widely expressed in the cortex—including the claustrum and piriform cortex—but are mostly absent from the basal ganglia and septum (Burnet et al., 1995). Cellular co-expression of D2 and 5-HT2A receptors is rare, but functional interactions between these receptors in the striatum may help reduce extrapyramidal side effects of D2 antagonists (Bubser et al., 2001). In our study, apart from the olfactory tubercle, the regions displaying the most extensive risperidone-induced transcriptional changes also exhibited low levels of D2 and 5-HT2A receptor expression—such as the septum, corpus callosum, and commissures—indicating that these effects are likely indirect. Similar results were recently observed with chronic olanzapine treatment (Abrantes et al., 2022), where differentially expressed genes were not enriched for dopaminergic pathways despite known interactions with D2-expressing neurons. Overall, these findings suggest that indirect mechanisms contribute to both acute and long-term transcriptional responses to antipsychotic medications.

We observed a robust and statistically significant overlap between risperidone-regulated genes and those linked to schizophrenia, based on non-targeted analyses using the largest available sets of gene-trait associations - GeDiPNet 2023 database and GWAS Catalog. This observation is consistent with recent reports that also pointed to antipsychotic drug-regulated genes as associated with the etiology of schizophrenia (Abrantes et al., 2022; Chestnykh et al., 2025). We demonstrate that the expression of schizophrenia-associated transcripts is not uniformly distributed across the forebrain. The majority of disease-related genes exhibit differential regulation of expression in the olfactory tubercle, lateral septal nucleus, and brain commissures, which together account for 15 of the 21 schizophrenia-linked genes. These results confirm previous findings and support a dopaminergic etiology of schizophrenia, even if part of the changes are indirect and observed outside the dopaminergic system. The common feature of these brain areas is a high content of glia, oligodendrocytes in particular. Notably, risperidone increased the expression of *Olig2* in the corpus callosum and anterior commissure (#6), which was previously found to have reduced expression in the brains of schizophrenic and bipolar patients (Tkachev et al., 2003). *Olig2,* as a key transcription factor, may affect the expression of other myelin-related genes, as it influences precursor and mature oligodendrocytes and is necessary and sufficient for the generation of oligodendrocytes and myelination (Georgieva et al., 2006). The downregulation of *Cacna1i* in the nucleus accumbens (#5) after risperidone administration is also of particular interest, given this region’s role in motivation, emotions, and limbic–motor integration (Salamone and Correa, 2012). *Cacna1i* encodes the Cav3.3 T-type calcium channel, which contributes to neuronal rhythmic firing. Considering that psychosis has been associated with sustained dopamine release and impaired habituation (Panayi et al., 2023), we speculate that reduced *Cacna1i* expression may attenuate phasic dopamine signaling and help mitigate hyperdopaminergic states. Notably, elevated *CACNA1I* expression has been reported in the hippocampus of schizophrenia patients (Xie et al., 2018), suggesting that risperidone may partially reverse disease-associated transcriptional changes. However, further studies, particularly in human tissue, are needed to confirm whether similar effects occur in the nucleus accumbens of patients treated with antipsychotics. Another important observation is the increased expression of *Smpd3* in the lateral septal nucleus (#13) following risperidone treatment. *Smpd3* encodes neutral sphingomyelinase-2 (NSM), an enzyme involved in sphingolipid metabolism and synaptic development. Abnormal sphingolipid metabolism has been implicated in schizophrenia and comorbid metabolic syndrome (Castillo et al., 2016; Chestnykh et al., 2025), though the specific role of *SMPD3* in cognitive decline and neurodegeneration remains unclear (Stoffel et al., 2018; Zhu et al., 2024). These findings further support the involvement of myelin dysfunction in schizophrenia (Takahashi et al., 2011) and link abnormal dopamine signaling with a dysfunction of oligodendrocytes. Finally, the enrichment revealed that schizophrenia-associated genes regulated by risperidone had increased frequency of binding sites for Polycomb group proteins. The involvement of SUZ12, MTF2, and CBX2 suggests that Polycomb-mediated chromatin remodeling may contribute to coordinated gene regulation across the forebrain. Functional annotations further indicated enrichment in processes related to synaptic transmission and neuronal morphology. Together, these findings point to upstream regulatory control and downstream functional convergence on pathways relevant to schizophrenia pathophysiology.

Our study has several limitations. First, the model involves the use of naïve animals and a single acute dose of 0.5 mg/kg risperidone. Arguably, the drug’s effects in a healthy animal may not accurately reflect the mechanism of action relevant to alleviating psychosis and other positive symptoms of schizophrenia. While this is undeniable, progressing to more complex models is not feasible until the baseline effects are established. We have previously demonstrated that the same model and drug dose are effective in analyzing the mechanisms of various psychotropic drugs (Korostynski et al., 2013). Moreover, it is debatable how well schizophrenia or its symptoms can be modeled in rodents (Powell and Miyakawa, 2006). New mouse models carrying mutations equivalent to rare human mutations associated with a very high risk of schizophrenia may provide a better approximation of the underlying pathology (Meechan et al., 2015; Rutkowski et al., 2021). However, analyzing differential gene expression with spatial resolution is a novel approach—the methodology is still evolving, and previous data, especially on the effects of acute antipsychotic treatment in naïve animals, have been largely unavailable. The present study aimed to fill this gap by offering one of the first brain-wide spatial transcriptomic datasets capturing gene expression responses to drug treatment. A second major limitation of our study stems from technical constraints of the method. Limited sequencing depth per tissue spot biases the dataset toward more abundant transcripts. The absence of *Fos* and several other activity-regulated transcripts may reflect sequencing limitations or the exclusion of low-abundance transcripts while handling a multidimensional, zero-inflated dataset. The spatial resolution is not at the single-cell level, resulting in a variable number of cells per spot, which further complicates interpretation. This is especially relevant for antipsychotics, as their primary target, the D2 receptors, are expressed in neurons within highly heterogeneous structures. For example, in the striatum, D2- and D1-expressing medium spiny neurons are distinct but notoriously difficult to separate, and often have opposing functions, such as mediating the effects of antipsychotics and psychostimulants (Bertran-Gonzalez et al., 2008). Therefore, a single spot in our dataset may contain a mixture of functionally different neurons, glia, and other cell types. This challenge may be addressed through methodological improvements and increased spatial resolution, although this would increase analytical complexity. While the observed expression patterns are unlikely to reflect the full range of transcriptional changes, they nevertheless enable region-specific assessment of risperidone effects and provide insight into the underlying mechanisms.

In summary, the spatial analysis of risperidone-regulated gene expression in the mouse forebrain offers significant new insights into the mechanisms of antipsychotic drug action. It shows a notable overrepresentation of schizophrenia-associated genes among differentially regulated transcripts, supports the idea of a dopaminergic mechanism in the disease’s etiology, and connects it to oligodendrocyte disruption based on spatial gene expression patterns. Further research is needed to confirm these mechanisms in models with strong construct validity and to enhance spatial resolution for single-cell transcriptome data.

## 5. Acknowledgments

The study was supported by the NCN Opus 20 2020/39/B/NZ7/01494 project and the statutory funds of the Maj Institute of Pharmacology of the Polish Academy of Sciences. The funders had no role in the study design, data collection and analysis, decision to publish, or preparation of the manuscript.

## 6. CRediT authorship contribution statement

*Magdalena Ziemiańska*: Investigation, Visualization, Formal Analysis, Validation, Writing – Original Draft Preparation, Writing – review & editing. *Mateusz Zięba*: Formal analysis, Data curation, Software, Visualization. *Anna Radlicka-Borysewska:* Investigation, Writing – review & editing, *Łukasz Szumiec*: Investigation, Writing – review & editing. *Sławomir Gołda*: Investigation, Validation. *Małgorzata Borczyk*: Methodology, Writing – review & editing. *Marcin Piechota*: Methodology, Formal analysis, Data curation. *Michał Korostyński*: Conceptualization, Supervision, Resources, Writing – review & editing. *Jan Rodriguez Parkitna*: Conceptualization, Funding Acquisition, Supervision, Resources, Project administration, Writing – Review & Editing.

---

**Supplementary Figure 1.**
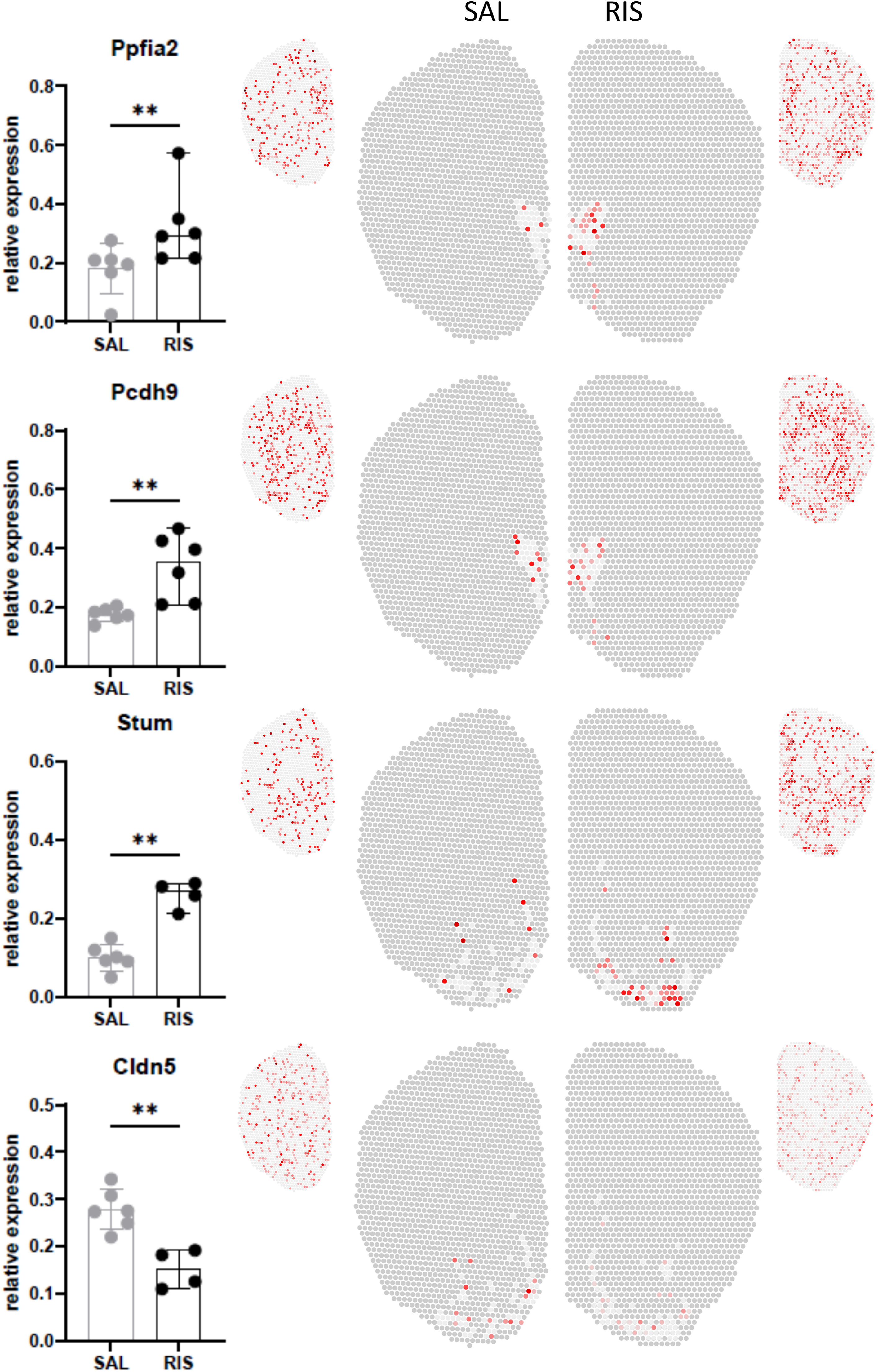
Additional spatial gene expression profiles of risperidone-regulated transcripts following acute risperidone treatment. The bar plots (left) and spatial gene expression profiles (right) illustrate the relative expression levels of genes regulated by risperidone and associated with schizophrenia in the previous human studies. Bar plots show medians with 95% confidence intervals, based on Mann–Whitney U test results (p1<10.01). Full expression profiles are displayed alongside the main images. Statistically significant changes were observed only in specific clusters, as outlined in the main panels.

**Supplementary Table 1.**
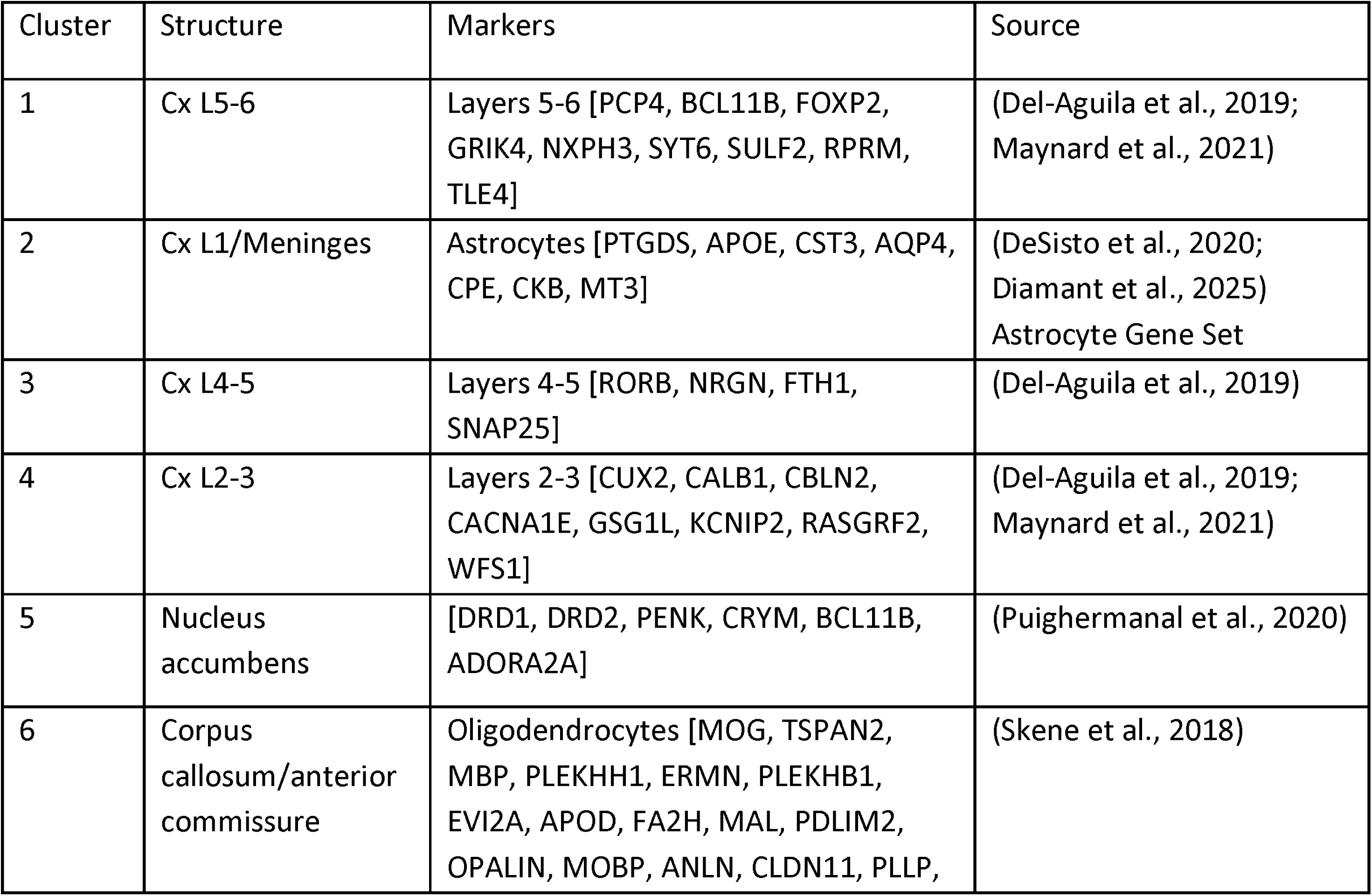
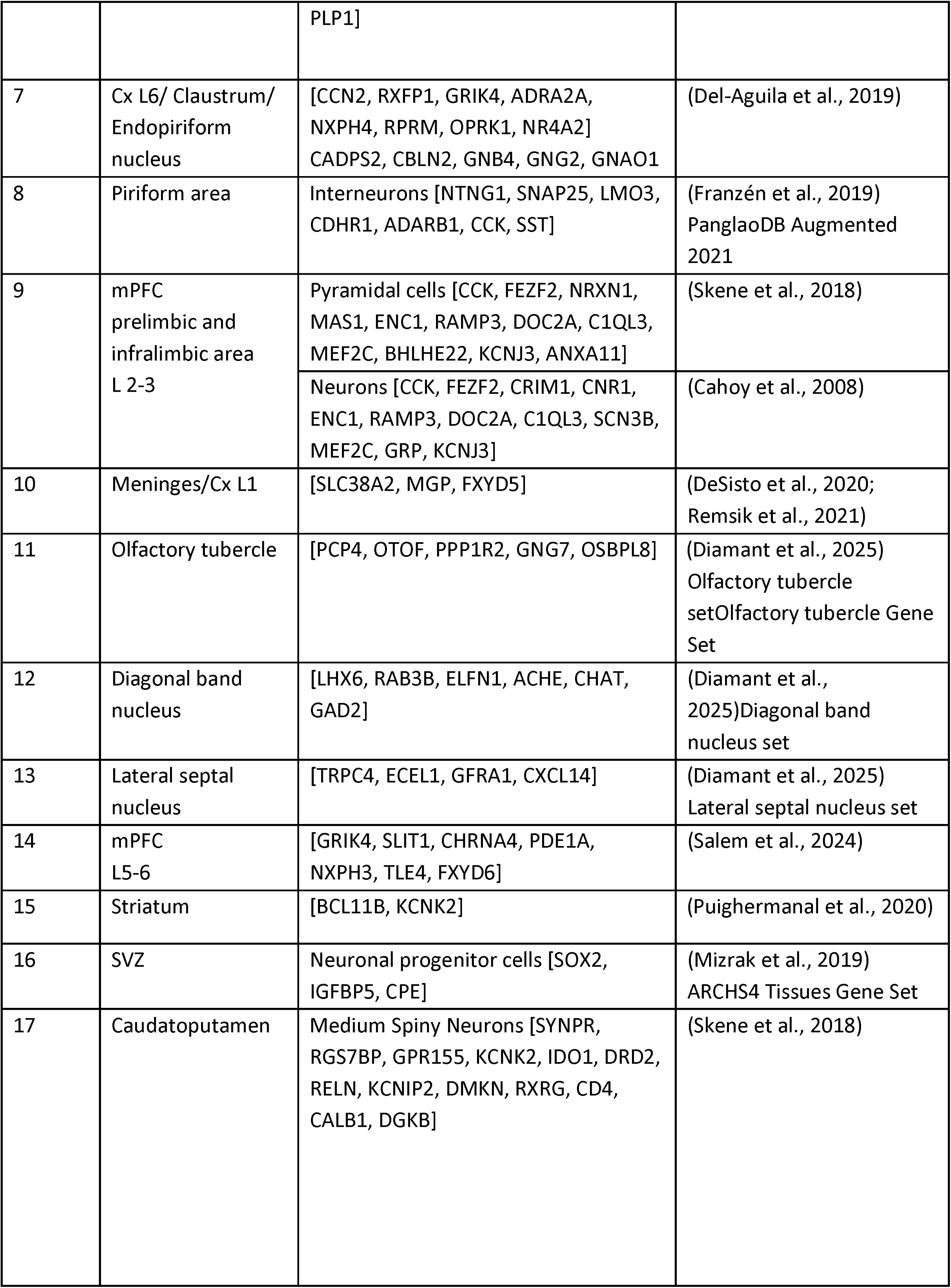
Identification of structural and cell-type markers based on literature and database sources.

**Supplementary Table 2.** Results of enrichment analysis for disease-related genes in the GeDiPNet 2023 database created with the Enrichr Appyter. The q-value is an adjusted p-value calculated using the Benjamini-Hochberg method for correction for multiple hypothesis testing.

| Term | p-value | q-value | overlap genes |
| --- | --- | --- | --- |
| Schizophrenia | 0.000171 | 0.056766 | EGR3, HOMER1, EGR4, LPAR1, PLEKHA6, OLIG2, CTCF, KALRN, LEMD2, SMPD3, CLDN5, CACNA1I, SYT11, BHLHE40, PHACTR3, NCAM1, SLC17A7, CALN1 |
| Atrophy of the Spinal Cord | 0.000462 | 0.076398 | FA2H, CCT5 |
| Hypocholesterolemia | 0.001194 | 0.131751 | UBE3B, CCT5 |
| Clonic Seizures | 0.002351 | 0.141523 | KCNA1, SLC2A1, SLC30A1, SLC17A7 |
| Spastic Paraplegia | 0.002501 | 0.141523 | FA2H, SLC2A1, DDHD1, CCT5 |
| Rubral Tremor | 0.002565 | 0.141523 | EGR3, SLC2A1 |
| Hypotonic Seizures | 0.004254 | 0.197108 | KCNA1, SLC2A1, SLC30A1, SLC17A7 |
| Dyskinesia | 0.005757 | 0.197108 | HOMER1, SLC2A1 |
| Focal Seizures | 0.006238 | 0.197108 | FA2H, KCNA1 |

**Supplementary Table 3.** List of terms connected with risperidone-regulated, schizophrenia-associated 21 genes obtained by meta-analysis of these genes in Enrichr-KG tool using the following databases: ChEA3 2022, MGI Mammalian Phenotype Level 4 2021, GO Biological Processes 2021, and Reactome 2022.

| database | full term | abbreviated term in Figure 4. |
| --- | --- | --- |
| GO Biological Processes 2021 | regulation of postsynapse organization (GO:0099175) | postsynapse organization |
| GO Biological Processes 2021 | cell chemotaxis (GO:0060326) | cell chemotaxis |
| GO Biological Processes 2021 | regulation of calcium ion-dependent | Ca<sup>2+</sup>-ion-dependent |
|  | exocytosis (GO:0017158) | exocytosis |
| GO Biological Processes 2021 | protein localization to chromosome (GO:0034502) | protein localization to chr |
| GO Biological Processes 2021 | neurotransmitter transport (GO:0006836) | neurotransmitter transport |
| ChEA3 2022 | RARB 27405468 Chip-Seq BRAIN Mouse | RARB |
| ChEA3 2022 | MTF2 20144788 ChIP-Seq MESC's Mouse | MTF2 |
| ChEA3 2022 | SUZ12 18692474 ChIP-Seq MESC's Mouse | SUZ12 PMID 18692474 |
| ChEA3 2022 | CBX2 27304074 Chip-Seq ESC's Mouse | CBX2 |
| ChEA3 2022 | SUZ12 20075857 ChIP-Seq MESC's Mouse | SUZ12 PMID 20075857 |
| Reactome 2022 | Protein-protein Interactions At Synapses R-HSA-6794362 | PPI at synapses |
| Reactome 2022 | Glutamate Neurotransmitter Release Cycle R-HSA-210500 | Glutamate release cycle |
| Reactome 2022 | Neurotransmitter Release Cycle R-HSA-112310. | Neurotransmitter Release Cycle |
| Reactome 2022 | Nuclear Events (Kinase And Transcription Factor Activation) R-HSA-198725 | Kinase and TF activation |
| Reactome 2022 | NGF-stimulated Transcription R-HSA-9031628 | NGF-induced transcription |
| MGI Mammalian Phenotype Level 4 2021 | abnormal dorsal root ganglion morphology MP:0000961 | abnormal DRG morphology |
| MGI Mammalian Phenotype Level 4 2021 | abnormal dendrite morphology MP:0008143 | abnormal dendrite morphology |
| MGI Mammalian Phenotype Level 4 2021 | abnormal miniature excitatory postsynaptic currents MP:0004753 | abnormal mEPSCs |
| MGI Mammalian Phenotype Level 4 2021 | abnormal proprioceptive neuron morphology MP:0004297 | abnormal PNs morphology |
| MGI Mammalian Phenotype<br>Level 4 2021 | decreased body height MP:0001255 | decreased body height |

